# The transcription factor TCF21 is necessary for adoption of cell fates by Foxd1+ stromal progenitors during kidney development

**DOI:** 10.1101/2024.08.14.607910

**Authors:** Gal Finer, Mohammad D. Khan, Yalu Zhou, Gaurav Gadhvi, George S. Yacu, Joo-Seop Park, R. Ariel Gomez, Maria Luisa S. Sequeira-Lopez, Susan E. Quaggin, Deborah R. Winter

## Abstract

Normal kidney development requires the coordinated interactions between multiple progenitor cell lineages. Among these, Foxd1+ stromal progenitors are essential for nephrogenesis, giving rise to diverse cell types including the renal stroma, capsule, mesangial cells, renin cells, pericytes, and vascular smooth muscle cells (VSMCs). However, the molecular mechanisms governing their differentiation remain poorly understood. This study investigates the role of Tcf21, a mesoderm-specific bHLH transcription factor, in Foxd1+ cell fate determination.

Using single-cell RNA sequencing (scRNA-seq), we analyzed 32,461 GFP+ cells from embryonic day 14.5 (E14.5) *Foxd1^Cre/+^;Rosa26^mTmG^;Tcf21^f/f^*kidneys (*Tcf21-cKO*) and controls. Clustering identified a predominant stromal population, further divided into six subpopulations associated with healthy kidney development: nephrogenic zone-associated stroma, proliferating stroma, medullary/perivascular stroma, collecting duct-associated stroma, differentiating stroma, and ureteric stroma. Loss of Tcf21 resulted in marked depletion of medullary/perivascular stroma, collecting duct-associated stroma, proliferating stroma, and nephrogenic zone-associated stroma stromal subpopulations, confirmed by immunostaining, which revealed severe constriction of medullary and collecting duct stromal spaces.

Additionally, we identified a novel cluster unique to *Tcf21-cKO* kidneys, characterized by high expression of Endomucin (Emcn), a vascular endothelial marker. These cells spanned across pseudotime trajectories and were distributed broadly across the mutant kidney. The emergence of Emcn-expressing cells in *Tcf21-cKO* kidneys coincided with a reduction in Acta2-expressing medullary stromal cells, suggesting a population shift.

Our findings highlight the critical role of Tcf21 in directing Foxd1+ progenitor differentiation. Loss of Tcf21 disrupts stromal cell fates, leading to aberrant kidney development and providing new insights into the mechanisms underlying congenital kidney anomalies.

**TRANSLATIONAL STATEMENT:** This study reveals critical insights into kidney development and congenital anomalies by identifying the developmental origins of stromal heterogeneity and the key role of Tcf21 in stromal progenitor differentiation. These findings enhance our understanding of stromal cell fate decisions and their relevance to congenital disorders. Additionally, this work provides valuable information for improving the recapitulation of the stromal compartment ex vivo, a current challenge in kidney organoid models. The role of Tcf21 in stromal phenotypic modulation underscores its broader significance in tissue repair and fibrotic diseases, suggesting potential avenues for therapeutic intervention.

## INTRODUCTION

Work that has spanned over seven decades increased our understanding of molecular mechanisms in kidney development and has been informing new regenerative based solutions for drug screening, disease modelling, and renal replacement ^1, 2^. The developing mammalian kidney is composed of four progenitor lineages: nephron (Six2+ cells), ureteric (Hoxb7+ cells), stromal/interstitial (Foxd1+ cells), and endothelial (c-kit+ cells) ^3, 4^. During kidney development, these progenitor populations signal to each other to ensure the ongoing expansion of the kidney to an organ of normal size and function ^5, 6^. The forkhead box D1 (Foxd1+) progenitors give rise to a diverse array of cell types including most of the renal stroma, vascular smooth muscle cells (VSMCs), pericytes, mesangial cells, renin-, and erythropoietin-producing cells ^7, 8^. Conventionally, renal stroma is divided into three anatomical regions: cortical, medullary, and papillary ^9^. Recent research, however, has identified a dozen distinct anatomical positions within the stroma in mice, each characterized by unique cell types, highlighting its heterogeneity ^10^. Foxd1+ cells play critical roles in branching morphogenesis, nephron development, and vascular patterning ^11^, possibly by promoting maturation of neighboring epithelial and endothelial cells ^12–15^. Despite these insights, understanding the molecular mechanisms through which the stroma influence non-stromal cells and guide the differentiation of Foxd1+ cells into distinct stromal populations is essential for normal kidney development, yet remains incomplete.

TCF21 is a mesoderm-specific transcription factor of the bHLH family that forms homo/heterodimers upon binding to its target DNA sequence ^16^. Originally named Pod1 ^17^, epicardin ^18^, or capsulin ^19, 20^, Tcf21 is expressed in the developing urogenital, cardiovascular, respiratory, and gastrointestinal systems during embryonic development. Tcf21 has been recently implicated as a causal gene for coronary artery disease in humans ^21, 22^. Within the kidney, Tcf21 has been implicated in the formation of the renal stroma and perivascular cells, influencing the structural integrity and function of the organ ^23^. To investigate the role of Tcf21 kidney stroma development and uncover the developmental origins of cell diversity arising from Foxd1+ cells, we performed scRNA-seq on normal and Tcf21-deficient Foxd1+ kidney cells from E14.5 mouse embryos, an early stage of nephrogenesis.

## MATERIALS AND METHODS

### Mouse strains and embryo staging

All mouse experiments were approved by the Animal Care Committee at the Center for Comparative Medicine of Northwestern University (Evanston, IL) and were performed in accordance with institutional guidelines and the NIH Guide for the Care and Use of Laboratory Animals. For gestational age, noon of the day on which the mating plug was observed was designated E0.5. Sex was determined before analysis and samples were pooled accordingly. The following mice were used in the studies described: *Tcf2*1*^floxf/^*^lox^ was created as previously described ^24^, *Foxd1-eGFPCre* (JAX Stock #012463), *Rosa26-mTmG* (JAX Stock: #007576). Genotyping was performed with Quick-Load Taq 2X Master Mix (M0271L, Biolabs) on 2% agarose gel for the PCR bands.

### Histology and immunohistochemistry

Mouse embryo dissections were performed at the indicated time-points. Samples were fixed with 10% Neutral buffered formalin or 4% paraformaldehyde overnight, embedded in paraffin, and sectioned into 5µm by the Mouse Histology and Phenotyping Laboratory of Northwestern University. Sections were deparaffinized in xylene and ethanol and boiled in citrate buffer for antigen retrieval. Sections were then treated with 1% donkey serum with 0.3% TritonX100 in PBS for 30 minutes at room temperature before incubating with primary antibodies at 4°C overnight. Primary antibodies: GFP (Abcam, ab13970, 1:500), Acta2 (Sigma, A2547, 1:400), Emcn V.7C7 clone (Abcam, ab106100, 1:250), and Cd31 (R&D, AF3628, 1:100). Secondary antibodies: AlexaFluor-488, 594, and 647 were incubated for 90 minutes at room temperature.

### Quantification and statistical analysis

Images were captured, and each channel was processed and thresholded to generate binary masks using FIJI/ImageJ software. The GFP+ EMCN+ CD31- area for stromal endomucin was quantified and normalized to the surface area of the kidney mid-section. Statistical analyses were performed using GraphPad Prism 10 (GraphPad Software, San Diego, CA). Group comparisons were conducted using an unpaired two-tailed t-test, with p-values <0.05 considered statistically significant.

### Preparation of single cells from E14.5 mouse embryos

Breeding triads were set up and allowed to conceive over a 24-hour period to ensure precise timing of pregnancy. E14.5 embryos from *Foxd1^Cre^;Rosa26^mTmG^;Tcf21^f/f^*and *Foxd1^Cre^;Rosa26^mTmG^;Tcf21^+/+^* were harvested from three dams on the same date. Kidneys were dissected under sterile conditions and washed in cold calcium- and magnesium-free PBS. To dissociate cells, a previously described protocol ^25^ was followed with modifications. Kidneys were minced and digested using a freshly prepared enzymatic mix containing Collagenase type 2 (2.5 mg/ml; Worthington, LS004174), DNase I (125 U/ml; Worthington, LS002058), and Bacillus licheniformis protease (7.5 mg/ml; Sigma, P5380). The samples were incubated at 14°C for 40 minutes with gentle pipetting every 5 minutes to aid dissociation. Digestion was halted using a stop solution comprising 20% FBS in calcium- and magnesium-free PBS. Cells were centrifuged at 1,250 rpm for 7 minutes to pellet them, then resuspended in 1 ml of DMEM (Dulbecco’s Modified Eagle Medium, Gibco, Netherlands) supplemented with 2% FBS. The suspension was passed through a 40-μm strainer (Scienceware Flowmi, H13680-0040) into DNA LoBind tubes (Eppendorf, 022431005). Following two additional filtrations into 5-ml polypropylene round-bottom tubes (Falcon™, 352063), the cells were resuspended in 500 μl DMEM with 2% FBS. Flow cytometry was performed at the Northwestern University Flow Cytometry Core using a BD FACSAria™ SORP system and a BD FACSymphony™ S6 SORP system. GFP-positive tdTomato-positive cells were sorted (**Suppl Fig. S1**), collected, and resuspended in PBS with 1% BSA (Bovine Serum Albumin) for immediate downstream applications.

Samples of the same genotype and sex were pooled to produce four final groups with the following total cell counts: Control Female: 199,522 cells (n=2), Control Male: 235,783 cells (from embryos n=4), Mutant Female: 109,099 cells (n=3), and Mutant Male: 178,086 cells (n=1). The average cell viability across all samples was greater than 86%, ensuring high-quality data for downstream analysis. For sequencing, 10,000 cells from each sample were uploaded, providing sufficient representation of the stromal populations for comparative transcriptomic profiling.

### Initial Processing of Single-cell RNA-seq

Single cells were processed using the 10x Genomics Chromium 3’ Kit version 3 and sequenced on an Illumina NovaSeq with a target of ∼400,000,000 reads per sample. Raw sequencing data was demultiplexed and processed using Cell Ranger v6.1.0 and a custom reference combining the mm10 transcriptome with the eGFP and tdTomato RNA sequences. Filtered count metrics from each sample were loaded as a Seurat object (v4.0.2). Genes were retained in the analysis only if expressed in at least three cells. Cells exhibiting fewer than 2000 or more than 8000 gene features were filtered out of the subsequent analysis. Moreover, cells with mitochondrial content exceeding 15% or with hemoglobin gene (Hba and Hbb) expression above 1%, were also discarded. Following this, Scrublet (v0.2.2) was used to identify cells likely to be doublets, applying 25 principal components for each sample. Across all samples, an expected doublet rate threshold of 7.6% (based on loading of approximately 16,000 cells) was set to ensure thorough and consistent doublet identification leading to 2701 cells marked for removal (**Suppl Fig. S2b**). After filtering, all samples were merged into a single Seurat object. Estimated cell numbers and initial quality control (QC) statistics are given in the table below:

| Sample | Pre-QC Cell # | Median Genes | Median UMIs | % Reads in Cell | Post-QC Cell # |
| --- | --- | --- | --- | --- | --- |
| Control<br>Female | 11,049 | 3,570 | 12,569 | 95.5 | 9526 |
| Control<br>Male | 11,995 | 3,946 | 15,391 | 94.7 | 9679 |
| Mutant<br>Female | 7,867 | 4,468 | 18,576 | 93.2 | 5830 |
| Mutant<br>Male | 9,543 | 4,387 | 19,294 | 94.0 | 7426 |
| **Total** |  |  |  |  | **32,461** |

### Cell Type Annotation

SCTransform was applied on 3000 variable genes for data normalization and scaling, with UMI count and percent mitochondria set as variables to regress. Principal component analysis (runPCA) was performed across all samples and the first 25 PCs were selected for Uniform Manifold Approximation and Projection (UMAP). Clustering (FindClusters) was also run with 25 PCs and a resolution parameter of 0.25. Clusters were optimized to capture the major cell types and were stable across a range of parameters. To identify *de novo* markers for the resulting 13 clusters, FindMarkers was run using a two-sided Wilcoxon rank-sum test (min. pct = 0.25) with adjusted p-value calculated using Benjamini-Hochberg method (**Suppl Table S1**). Individual clusters were grouped together based on annotated cell type for further analysis.

### Stromal Subclustering

To identify sub-populations within stromal clusters, we selected all cells from the clusters annotated as stroma (n = 22,355). Cells exhibiting more than 10% mitochondrial content were excluded to eliminate stressed or dying cells that could skew the analysis.

| Sample | All Stromal Cells | Median Genes | Median UMIs | Filtered Stromal Cells |
| --- | --- | --- | --- | --- |
| Control<br>Female | 8,912 | 3,577 | 12,558 | 8,897 |
| Control<br>Male | 6,724 | 3,889 | 14,559 | 6,689 |
| Mutant<br>Female | 4,939 | 4,668 | 19,993 | 4,934 |
| Mutant<br>Male | 1,780 | 4,046 | 14,959 | 1,777 |
| **Total** | **22,355** | **3,922** | **14,644** | **22,297** |

The remaining 22,297 cells were renormalized using scTransform with 3000 variable genes and regressing on the difference between S and G2M score (CellCycleScoring) as recommended by Seurat to limit the effect of cell cycle on clustering while maintaining differentiation-associated trends. UMAP and clustering (FindClusters) were performed with the first 15 principal components and resolution at 0.20 based on the elbow of the scree plot and robustness of clusters (**Suppl Fig. S6a-b**). *De novo* markers across stromal clusters were identified as described above for cell types (**Suppl Table S2a**). Transcriptional signatures of stromal compartments were calculated using AddModuleScore on curated markers based on publicly available sources ^10, 26^, Allen Developing Mouse Brain Atlas (https://developingmouse.brain-map.org/), GUDMAP^27, 28^, and our previous studies ^23^ (**Suppl Fig. S4**). To compare our annotations to previously published clusters (Combes et al), we calculated the fraction of the top 20 published markers that overlapped our list of *de novo* markers by cluster (**Suppl Table S2b**).

### GenePaint

The spatial expression of representative key genes in control E14.5 mouse kidney was examined using *GenePaint* gene expression atlas (https://gp3.mpg.de/).

### Comparison of Stromal Cells in *Tcf21*-*cKO* and Control

Before comparing stromal cells across genotype, each sample was subsampled to 1777 stromal cells to match the lowest sample and ensure even numbers across sexes. This approach was used to take advantage of the independent samples but control for the effect of sex. The scProportionTest algorithm (https://github.com/rpolicastro/scProportionTest) was used to calculate p-values for differences in composition between *Tcf21-cKO* and control conditions using a permutation test.

Confidence intervals were provided through bootstrapping. Differential expression analysis was performed using FindMarkers to measure changes in expression distribution across all single cells in the stromal compartment in *Tcf21-cKO* vs. control (min.pct = 0.25). All stromal cells were included to avoid biases due to uneven clusters between conditions while comparing single-cell level expression. Genes with log fold change ≥ 0.25 and with p-adjusted value less than ≤ 0.05 were defined as significantly increased or decreased in expression. Enriched GO terms for genes that exhibited significantly increased or decreased expression in *Tcf21-cKO* were determined using the R package clusterProfiler (https://github.com/YuLab-SMU/clusterProfiler) and compared all 28,891 genes expressed in the dataset as background.

### Trajectory Analysis

To construct a pseudotime trajectory representing healthy differentiation, cells from control samples (n = 15,579, excluding 7 outlier cells from the ECM-expressing cluster) were isolated from the stromal population. We excluded the *Tcf21-cKO* cells in an effort to capture the relationship between subpopulations in normal development. This subset of cells was normalized using scTransform with regression by cell cycle score difference as above (S-G2M). Then, RunPCA and RunUMAP (dim=15) was performed. The two-dimensional representations was converted to be compatible with Monocle v3 ^29, 30^. Clustering was performed in Monocle using the cluster_cells function with default parameters. To build the underlying trajectory, learn_graph was run with parameters close_loop=FALSE and ncenter=310 (chosen based on calculating 2% of the number of cells). Pseudotime for each cell was calculated using order_cells()with the starting node set as the nephrogenic zone cluster. The function graph_test() was used to identify co-regulated genes across pseudotime with the autocorrelation approach (“principal_graph” parameter). Significant genes were defined by q value < 0.05. Genes were grouped into trajectory-based modules using find_gene_modules() with default parameters testing a range of resolution values from 10^−6^ to 10^-1^ and the aggregate_gene_expression was used to calculate module expression. The pseudotime calculated in the control samples was binned into discrete intervals (bins) of 3 units each, ranging from 0 to 30. To transfer the pseudobins from the control sample to *Tcf21-cKO* samples, the FindTransferAnchors() and TransferData() functions were used to identify the anchors with shared annotations between control and the *Tcf21-cKO* cells using the PCA as a dimensionality reduction technique function. To calculate enrichment of each trajectory-based module in the Emcn- expressing cluster in *Tcf21-cKO* samples, the percent of cells with score greater than 0 in this cluster compared with the rest of the sample was calculated. These numbers were used in a hypergeometric test to determine the p-value for enrichment.

## RESULTS

### Single-cell profiling of E14.5 *Foxd1* GFP+ cells identifies diverse cell populations within the developing kidney

To investigate the role of Tcf21 in the Foxd1+ stromal progenitors and their derivative cells at a single-cell level, we purified GFP-positive tdTomato-positive cells from male and female *Foxd1^Cre/+^;Rosa26^mTmG^;Tcf21^f/f^*(*Tcf21*-cKO) and littermate controls *Foxd1^Cre/+^;Rosa26^mTmG^;Tcf21^+/+^*via flow cytometry. In this model, *Tcf21* gene is permanently deleted, tdTomato expression is terminated, and GFP expression is maintained in all the progeny of Foxd1-expressing cells. We obtained 13,256 cKO and 19,205 control cells after QC (**Suppl Fig. S2a**). We defined 13 clusters using an unsupervised approach (see Methods) and assigned cell lineage annotations based on *de novo* markers and canonical gene expression (**Fig.1a-b, Suppl Fig. S2cd, Suppl Table S1**). This analysis identified a large population of stromal cells (Foxd1+ Meis1+ Pdgfra+) as well as representation of the nephron progenitor cell (NPC) lineage (Six2, Cited1, Crym), and significantly smaller populations of ureteric epithelium (Ret, Wnt 11, Gata3), immune cells (Fcer1g, Lyz2), and neurons cells (Tubb3, Map2) (**Fig. 1a-b, Suppl Fig. S2d-e**). Cycling genes were expressed across cell lineages (**Suppl Fig. S2f**). We confirmed that GFP expression was robust in both stromal and non-stromal cells, though Foxd1 expression was largely limited to the stromal lineage (**Fig. 1c**). Low expression of tdTomato and depleted expression of Tcf21 in *Tcf21*-*cKO* cells validate the activity of cre recombinase in these cells (**Fig. 1c, Suppl Fig. S2g**). Non-stromal GFP+ cells likely represent early transient expression of Foxd1 before the onset of lineage boundaries. All cell types were represented in both males and females of *Tcf21-cKO* and control samples, however the number and proportion of stromal cells was lower in sex-matched mutant samples (**Fig. 1d**, **Suppl Fig. S2h**). These differences in cell numbers are likely attributable to experimental variation rather than being biologically driven.

**Figure 1.**
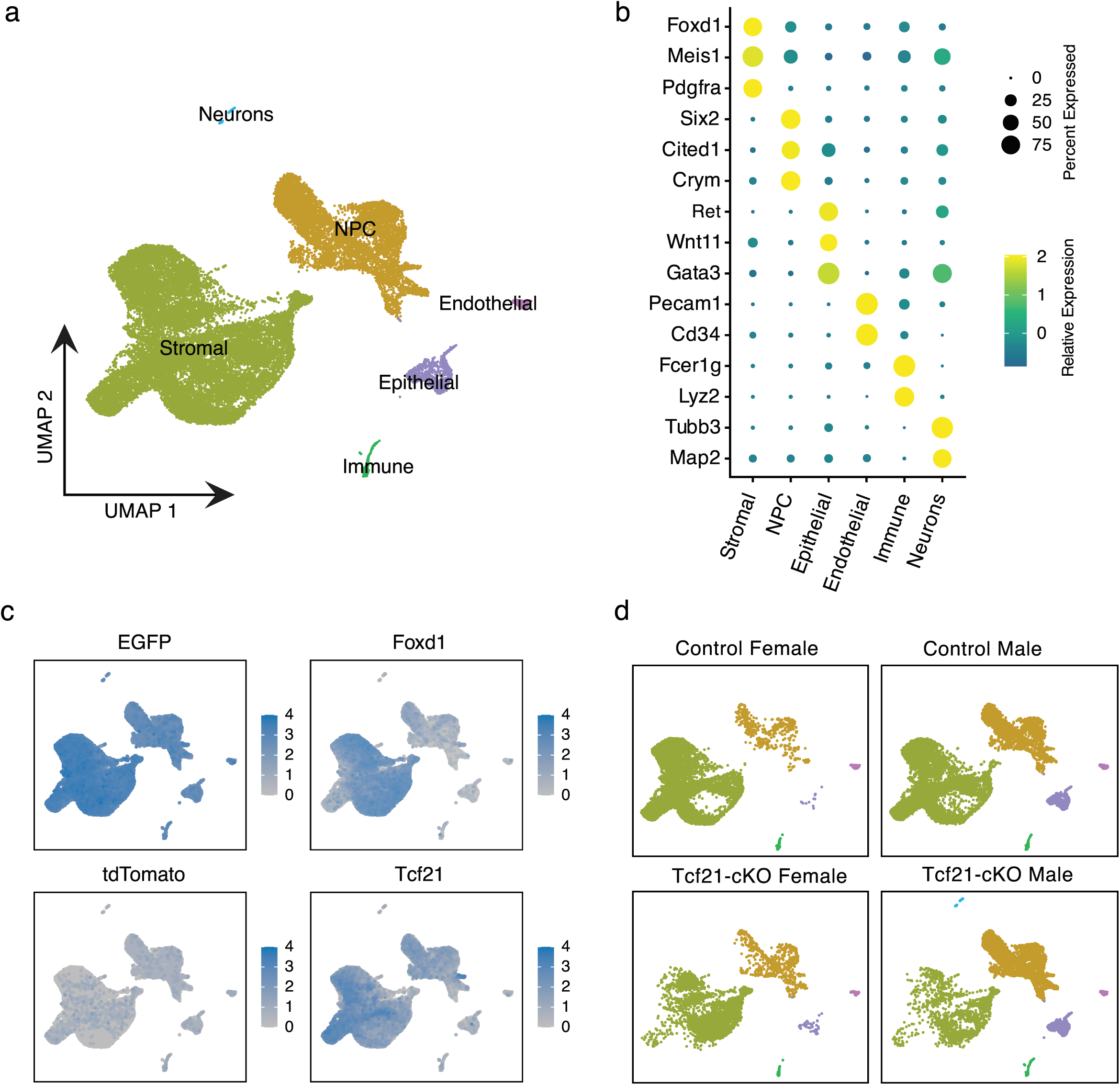
scRNA-seq analysis of the Foxd1-GFP+ tdTomato+ sorted cells from E14.5 embryonic mouse kidneys of Tcf21-conditional knockout (*Tcf21-cKO*) and controls. (a) UMAP plot of 32,461 individual cells combined from male and female *Tcf21-cKO* and control kidneys coloured by cell type annotation. (b) DotPlot illustrating expression of canonical genes across cell types. Size of dot illustrates percent of cells expressing the gene and colour scale indicates relative expression across cell types. (c) Feature plots of EGFP, Foxd1, tdTomato, and Tcf21 gene expression (log_2_ transformed). (d) UMAP plot of samples by experimental group and sex. See **Suppl Fig. S2h** for number of cells per experimental group.

### E14.5 stromal progenitors express genes associated with different cell fates

Next, we sought to explore the heterogeneity within the developing stromal cell populations by sub-clustering cells annotated as stroma (**Suppl Fig. S3a-b**). We defined seven stromal subpopulations (numbered 0-6) in *Tcf21*-*cKO* and controls with distinct gene expression profiles based on *de novo* marker expression (**Fig. 2a-b**). All clusters expressed similar levels of GFP and low level expression of tdTomato although expression of *Foxd1* and *Tcf21* was variable (**Suppl Fig. S3c**). To annotate these clusters, we created modules associated with different subpopulations based on canonical markers from the literature (**Suppl Fig. S4**) and visualized their expression along with canonical cell cycles genes (**Fig. 2c**). Based on module expression and *de novo* genes markers, we annotated Nephrogenic zone associated stroma; Proliferating stroma; Medullary/Perivascular associated stroma; Collecting duct associated stroma; Differentiating stroma; and Ureteric stroma. To confirm our annotations, we compared expression of key genes in our single-cell data vs. spatial expression from the *GenePaint* gene expression atlas (**Fig. 3**). Our annotations largely agreed with previously published stromal clusters during kidney development (Combes et al ^26^), with subtle differences (**Suppl Fig. S3d-e**). Our annotated Differentiating stroma cluster (cluster 3) more closely resembles the published Ureteric cluster. However, the lower levels of *Tbx18*, a gene know to play a critical role in early separation of the ureteric mesenchymal lineage from other mesenchymal lineages ^11^ ^31^, in these cells compared with those in cluster 6 did not support its classification as Ureteric. Moreover, *de novo* markers with unique expression in cluster 6 demonstrated high spatial specificity to the ureteric zone (**Fig. 3**). Instead, cluster 3 was annotated as "Differentiating Stroma" due to more diffuse expression of associated markers (**Fig. 3)**, consistent with its lack of a distinct anatomical location (**Fig. 3b**). Additionally, we annotated cluster 2 as Emcn-expressing based on unique expression of the gene Endomucin (*Emcn*), further discussed in the following section (**Fig. 2, Suppl Fig. S3f**).

**Figure 2.**
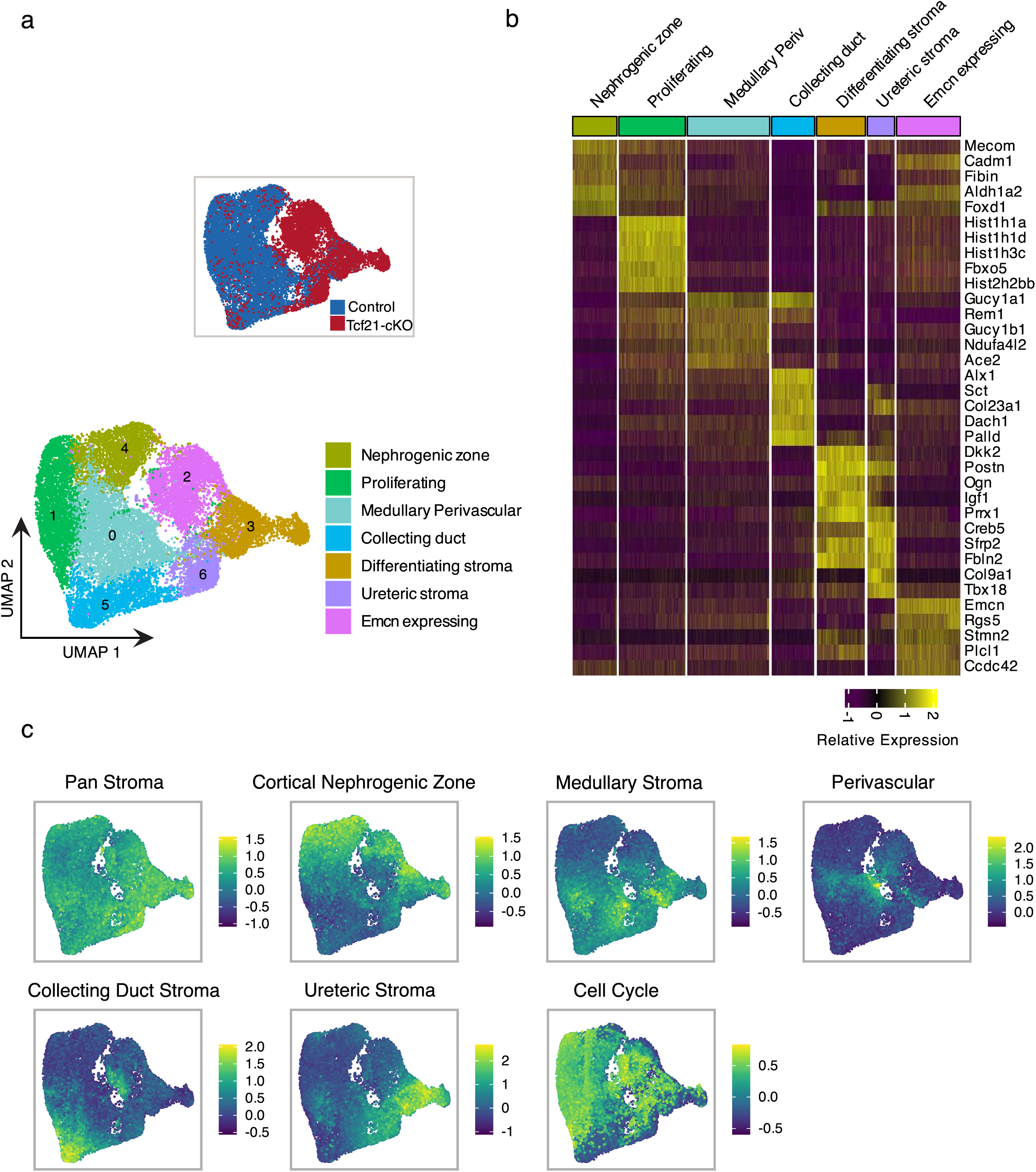
Stromal lineage sub-clustering identifies populations of distinct cell fate within the developing mouse kidney. (a) Sub-clustering of stromal cells (n=22,297) from E14.5 Foxd1GFP+ kidney cells (stromal cluster in **Fig. 1a**) identifies seven transcriptionally distinct populations and their suggested annotation. (b) Heatmap depicting the relative expression of 5 of the top de novo marker genes for each cluster across individual cells. Column color coding is consistent with **Fig. 2a**. (c) Feature plot of the module score distribution in UMAP space based on canonical gene signatures (listed in **Suppl Fig. S4**).

**Figure 3.**
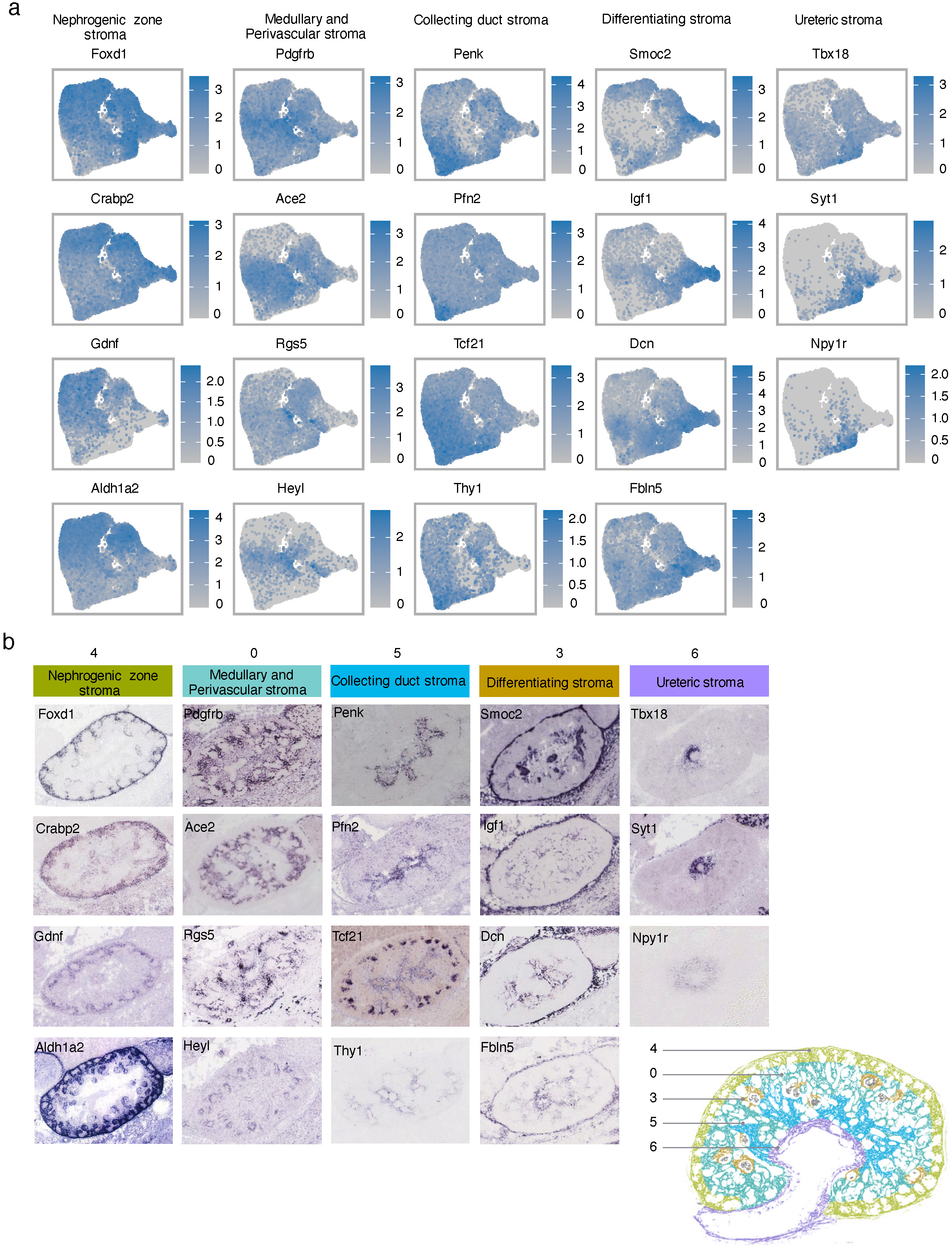
Spatial annotation of stromal clusters of control and *Tcf21*-*cKO*. (a) Featureplot showing the log_2_-transformed gene expression of markers for Nephrogenic zone stroma (*Foxd1, Crabp2, Gdnf, Aldh1a2*), Medullary and Perivascular stroma (*Pdgfrb, Ace2, Rgs5, Heyl*), Collecting duct stroma (*Penk, Pfn2, Tcf21, Thy1)*, Differentiating stroma (*Smoc2, Igf1, Dcn, Fbln5*), and Ureteric stroma (*Tbx18, Syt1, Npy1r)*. (b) Expression of the same marker genes by *in situ* hybridization in control E14 mouse kidneys as per *GenePaint* gene expression atlas (https://gp3.mpg.de/, cropped images from whole mount), includes schematic illustration of the spatial location of each stromal population in the developing kidney. Cluster 0 (Medullary/Perivascular stroma), cluster 4 (Nephrogenic zone stroma), cluster 5 (Collecting duct stroma), cluster 3 (Differentiating stroma), and cluster 6 (Ureteric stroma).

### Tcf21 is required for emergence of stromal sub-populations during kidney development

Next, we sought to determine how stromal progenitor cells in *Tcf21*-*cKO* have deviated from controls using a downsampled dataset to control for sex and cell number (see Methods). Strikingly, medullary/perivascular stroma, collecting duct-associated stroma, proliferating stroma, and nephrogenic zone-associated stroma clusters were dramatically reduced in *Tcf21-cKO* compared to control (**Fig. 4a-c**). Conversely, the numbers of differentiating stroma and ureteric-associated stroma clusters were moderately increased in *Tcf21-cKO*, though this may be due to reduced total stromal cell numbers. Notably, the most profound change was that the Emcn-expressing cluster was virtually exclusive to *Tcf21-cKO* kidneys, with only few cells (7 cells) of shared transcriptional profile identified in stromal cells of controls (**Fig. 4a-c**). These results were conserved when male and female samples were compared independently (**Suppl Fig. S6c-e**).

**Figure 4.**
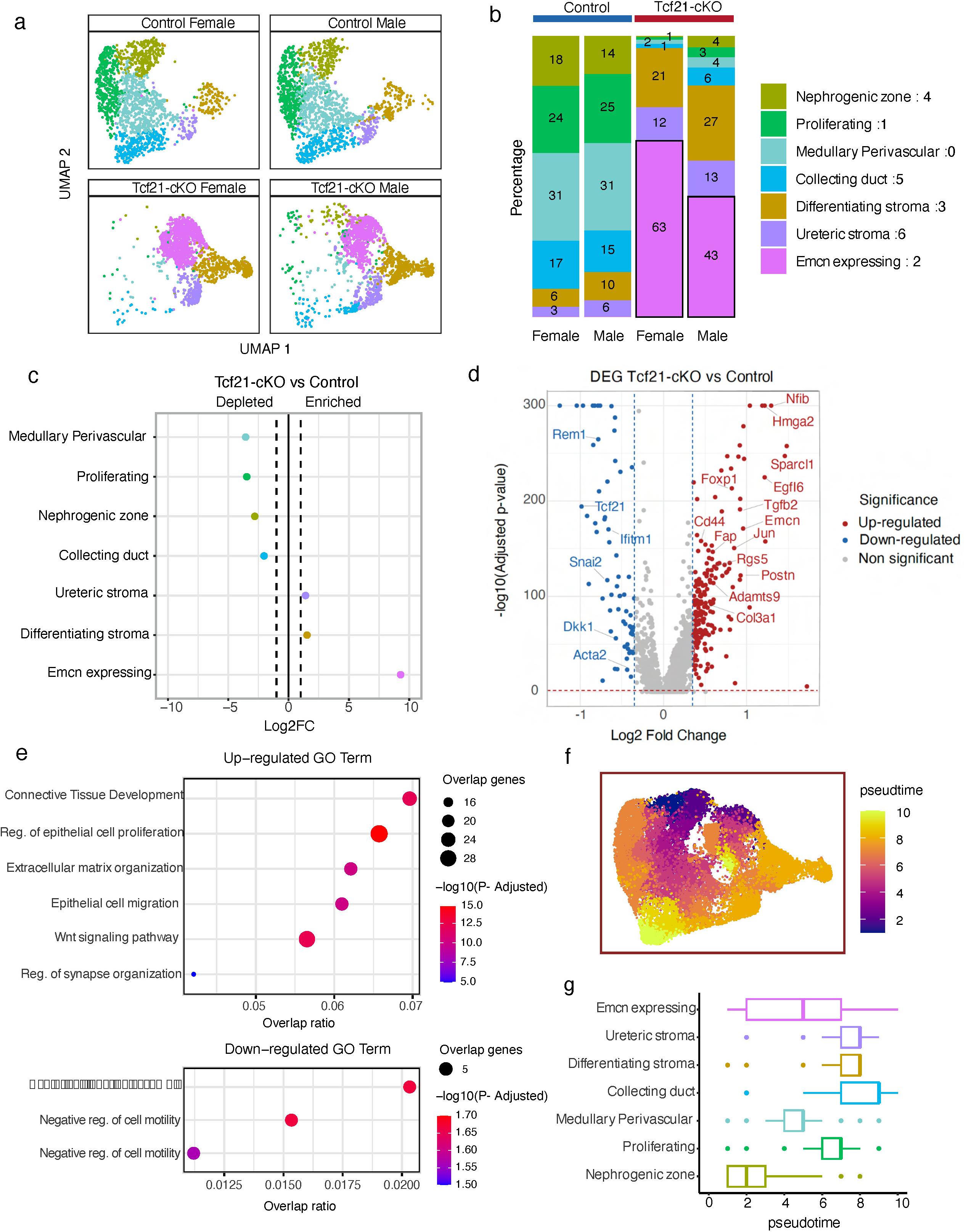
Tcf21 is critical for the development of specific stromal cell fates. (a) UMAP of Foxd1GFP+ 7,108 stromal cells (downsampled to equally reflect 4 samples) from male and female E14.5 control and *Tcf21-cKO* mouse kidneys (*Foxd1^Cre^;Rosa26^mTmG^;Tcf21^+/+^*and *Foxd1^Cre^;Rosa26^mTmG^;Tcf21^f/f^*, respectively). (b) Stacked bar chart indicating subpopulation composition by sample. Percent of each subpopulation in a sample are given. (c) Relative differences in cell proportions for each cluster between the *Tcf21-cKOs* versus controls. Dashes vertical lines mark absolute Log_2_ fold change (FC) > 1 compared with the control. All changes in proportion were significant after FDR was applied (scproportion permutation test; *n* = 3554/group; **Suppl Table S4a**). Bi-sex comparison in **Suppl Fig. S6c-e**. (d) Volcano plot of differentially expressed genes (DEGs) between all stromal cells in *Tcf21-cKOs* versus controls defined by log_2_(fold change) >0.25 and adjusted p-value <0.05. Red and blue dots denote significantly up- and down-regulated differential genes, respectively. (e) DotPlot of GO Terms significantly enriched among up-regulated (top panel) and down-regulated (bottom panel) DEGs. X axis denotes the overlap ratio (# genes in overlap/# genes in GO term, size of dot denotes the #genes in overlap, and color scales deotes the -log_10_ adjusted p-value (p-adj). (f) Feature plot of stromal cells with color-coding based on pseudotime bins. g) Box and whiskers plot depicting the range of pseudotime by bin in each stromal cluster.

In parallel, we performed differential gene expression analysis of the whole stromal compartment between *Tcf21-cKO* and control (**Suppl Table S3b**). We identified 214 genes up-regulated and 64 genes down-regulated in *Tcf21-cKO* compared with control cells (**Fig. 4d**). Tcf21 is among the most down-regulated genes, as expected, while genes associated with connective tissue development, epithelial cell proliferation, extracellular matrix organization, and WNT pathway were upregulated (**Fig. 4e**). In order to determine how the Emcn-expressing cells fit into healthy stromal cell differentiation, we ran Monocle on control stromal cells to estimate the underlying pseudotime trajectory assuming nephrogenic zone-associated cells represented the most progenitor-like subpopulation. By transferring control-based pseudotime to *Tcf21-cKO* cells, we found that the Emcn-expressing cluster comprised cells spanning across all pseudotime (**Fig. 4f-g**). Moreover, when we defined modules of co-regulated genes along the control trajectory, we found that specific cell subsets within the Emcn-expressing cluster expressed genes associated with a variety of other subpopulations including nephrogenic (module 18), proliferating (module 20), and medullary/perivascular (module 14) (**Suppl Fig. S5c-d**). Taken together, these results suggest that the *Tcf21-cKO* is associated with the emergence of cells displaying altered differentiation outcomes, including a potential aberrant differentiation.

To spatially define the defect in the stromal lineage of the *Tcf21-cKO*, we performed lineage-tracing of Foxd1 (GFP+) in *Foxd1^Cre/+^;Rosa26^mTmG^;Tcf21^f/f^* and control mice from E14.5 to E18.5. *Tcf21*-cKO demonstrated a marked constriction of the medullary stroma and the stromal space adjacent to the collecting ducts compared with controls (**Fig. 5a**). Additionally, immunostaining for endomucin, a marker of the stromal population exclusively expressed in *Tcf21-cKO*, confirmed robust expression solely in the stromal lineage of *Tcf21-cKO* kidneys at E14.5 and E16.5 (**Fig. 5b-c**). Furthermore, staining with alpha smooth muscle actin (Acta2), which marks medullary stromal cells ^10, 23^, demonstrated a loss of Acta2-expressing stroma and the emergence of Emcn-positive stromal cells in *Tcf21-cKO* kidneys (**Fig. 5d**), supporting a population shift. Taken together, these data support the scRNA-seq analysis results above and indicate a requirement for Tcf21 in the emergence of specific derivatives of the stromal lineage during kidney development.

**Figure 5.**
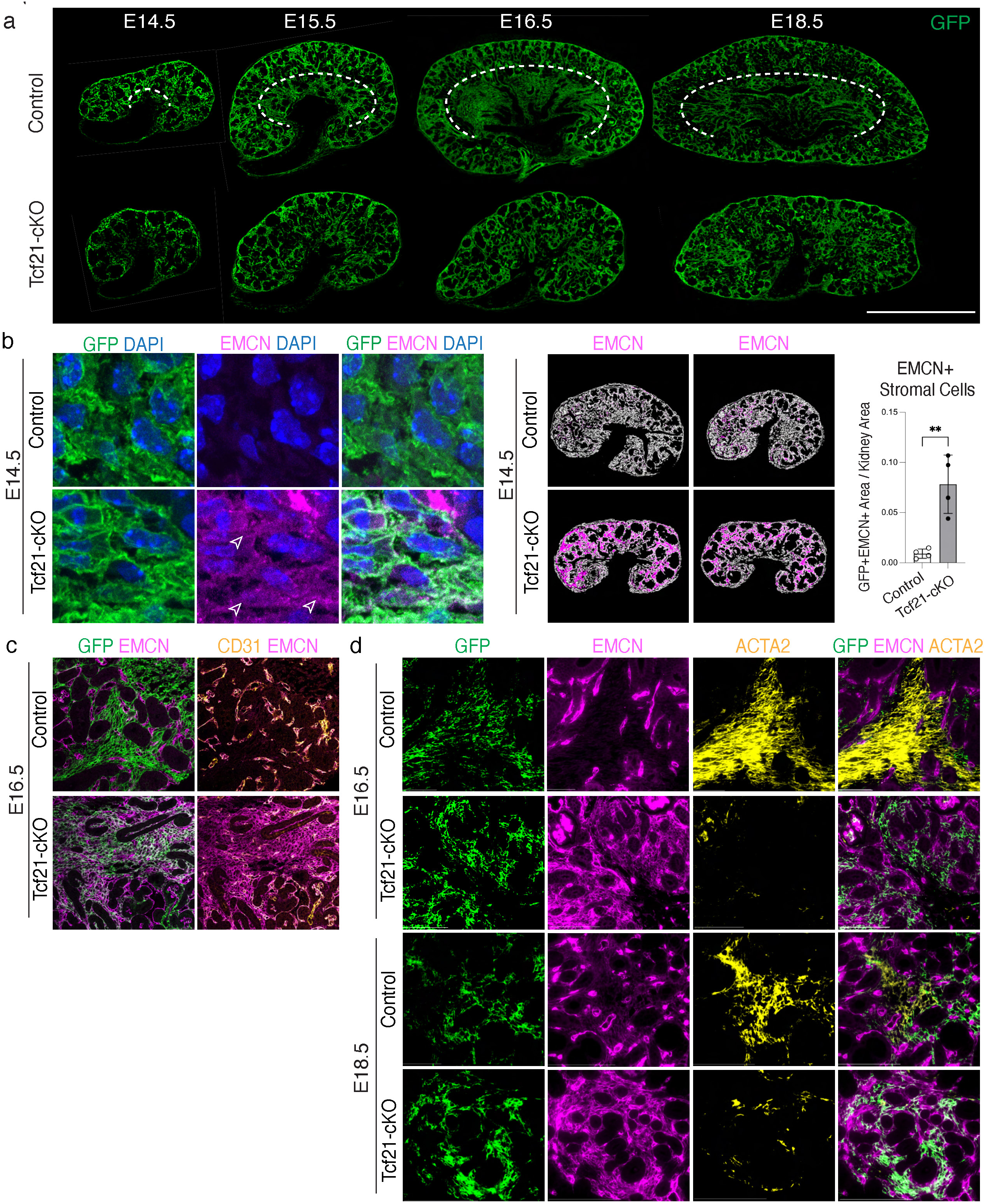
Lineage tracing and population shift in Foxd1+ stromal cells of *Tcf21-cKO* kidneys. Immunofluorescence staining of *Foxd1^Cre^;Rosa26^mTmG^;Tcf21^f/f^* and control kidneys. (a) Staining with an anti-GFP antibody (marking Foxd1+ lineage) shows a significant reduction in stromal space in *Tcf21-cKO* at E14.5, consistent with scRNA-seq findings. Lineage tracing at E15.5, E16.5, and E18.5 indicates that this reduction persists in medullary stroma and stroma adjacent to the collecting ducts in *Tcf21-cKO* kidneys (dotted line). Scale bar 500 μm. (b) Staining with an anti-endomucin (Emcn) antibody confirms the presence of novel Endomucin-positive stromal cells (arrowheads) in *Tcf21-cKO* at E14.5, occupying the majority of the stromal space (quantification of Emcn-positive stromal cells on the right, *p*=0.003). (c) Emcn- expressing stromal cells are again identified in the *Tcf21-cKO* kidney at E16.5. In contrast, in control kidneys at E16.5, Endomucin expression is restricted to the endothelium (co-expressed with CD31) and is absent in stromal cells. (d) Staining with an anti-Acta2 (alpha smooth muscle actin) antibody shows a severe reduction of Acta2-positive stromal cells *Tcf21- cKO*, which coincides with the emergence of Emcn-positive stromal cell at E16.5 and E18.5. Total number of embryos analyzed: n = 20 (4 male controls, 2 female controls, 5 male mutants, 1 female mutant, 4 controls with sex not determined, and 4 mutants with sex not determined).

## DISCUSSION

Our findings underscore the critical role of Tcf21 in the differentiation and fate determination of Foxd1+ stromal progenitors during kidney development. Using scRNA-seq and detailed sub-clustering analysis, we identified seven distinct stromal populations within the developing kidney. The deletion of Tcf21 led to a significant reduction in specific stromal populations, including medullary/perivascular stroma, collecting duct-associated stroma, proliferating stroma, and nephrogenic zone-associated stroma. Concurrently, a novel stromal population emerged in Tcf21-deficient kidneys, characterized by high expression of Endomucin (Emcn), a sialomucin family gene typically expressed in vascular endothelial cells. This novel stromal population was annotated as Endomucin-expressing stroma.

The kidney stromal compartment is known for its high heterogeneity, with Foxd1+ progenitors giving rise to diverse cell types, including fibroblasts, VSMCs, pericytes, renin cells, and mesangial cells ^7^. Previous studies have highlighted the pivotal roles of Foxd1+ cells in branching morphogenesis, nephron development, and vascular patterning^10, 32^. However, the molecular mechanisms guiding the differentiation of these cells into distinct stromal lineages have remained poorly understood.

The medullary and collecting duct stromal compartments are critical for the kidney’s structural integrity and function. The medullary stroma/interstitium maintains the osmotic gradient required for urine concentration, while the collecting duct-associated stroma supports collecting duct development and function. Previous studies ^23^ demonstrated the broad role of Tcf21 in medullary stroma formation and highlighted urinary concentration defects in mutants. However, they did not provide mechanistic insights into the developmental origins of stromal cell heterogeneity.

The identification of a distinct stromal population exclusively present in Tcf21-depleted Foxd1+ cells, characterized by unique Emcn expression, represents a significant finding. Pseudotime analysis revealed that the Emcn-expressing population is distinct within the stromal trajectory (**Fig. 4g**), diverging from the primary differentiation pathway of nephrogenic stroma. This divergence raises questions about their developmental state, including the possibilities that these cells are arrested at a specific stage, exhibit premature differentiation, undergo transdifferentiation, or display phenotypic modulation. Phenotypic modulation may enable these cells to retain certain stromal features while expressing markers typically associated with an alternate lineage. Histological analysis confirmed the widespread presence of this novel Emcn- expressing stromal population in the *Tcf21-cKO*, with endomucin-positive cells distributed broadly across the *Tcf21-cKO* kidney (**Fig. 5b-c)**. Interestingly, the emergence of Emcn-expressing cells coincided with a marked reduction in Acta2- expressing medullary stromal cells, suggesting a population shift. In the *Tcf21-cKO*, Emcn was expressed in cells previously positive for Acta2 (**Fig. 5d**), supporting phenotypic modulation and emphasizing the role of Tcf21 in maintaining stromal cell identity.

Interestingly, an increase in ureteric-associated stroma and differentiating stroma populations was observed in the Tcf21- cKO kidneys, though this may be due to reduced total stromal cell numbers. This expansion may represent a compensatory response to the depletion of other stromal populations, potentially driven by homeostatic imbalance and activation of signaling pathways aimed at preserving tissue structure and function. Future experiments, such as lineage tracing, could clarify whether these cells proliferate directly in response to the loss of other cell types.

Gene Ontology enrichment analysis revealed significant changes in genes associated with connective tissue development, extracellular matric organization, epithelial cell proliferation, and WNT signaling pathways in *Tcf21-cKO* stromal cells compared to controls.

Our findings highlight Tcf21’s pivotal role in Foxd1+ stromal progenitor differentiation, regulating distinct stromal populations for proper kidney development and function. These results also suggest broader implications for Tcf21 in mesodermal development and its relevance to fibrotic diseases with excessive ECM deposition.

This study has several constraints that should be acknowledged. The use of single-cell RNA-seq to understand Tcf21’s role in kidney development is challenged by the inclusion of GFP+ non-stromal cells in our samples, which may result from leakiness in the Cre-lox system, transient Foxd1 expression, or autofluorescence overlap with the GFP channel. Foxd1 expression in nephron progenitor cells (NPCs) has been reported by others ^8, 33–36^, likely due to a shared signature during early nephrogenesis ^26^. Although potential disruption of normal Foxd1 expression by the Cre model remains a consideration, pooling mice and including Foxd1-Cre controls mitigates this issue.

The limited cell numbers obtained from single embryos necessitated pooling, and uneven representation of sexes in biological replicates, influenced by Mendelian distribution and the requirement to harvest kidneys in a single experiment to avoid batch effects, posed additional challenges. Nonetheless, both male and female samples were included, with consistent findings across sexes, and immunostaining validation with a minimum of three replicates per condition addressed this issue. Finally, reliance on a single time point for scRNA-seq limits the ability to fully annotate cell fates and interpret differentiation trajectories. Future studies incorporating multiple time points will be essential to explore the transcriptional networks, direct targets, and signaling pathways regulated by Tcf21 in Foxd1+ cells.

## DATA AVAILABILITY

Data will be uploaded to the Gene Expression Omnibus upon acceptance of the manuscript.

## SUPPLEMENTAL MATERIAL

Suppl Fig. S1: Flow cytometry analysis of single-cell suspensions from E14.5 mouse kidneys for scRNA-seq.

Suppl Fig. S2: Single-cell RNA-seq analysis of Foxd1GFP+ sorted cells from control and *Tcf21-cKO* kidneys.

Suppl Fig. S3: Single-cell RNA-seq analysis of stromal cells

Suppl Fig. S4: Stromal subpopulation markers for module scores

Suppl Fig. S5: Comparison of stromal cells between *Tcf21-cKO* and controls

Suppl Fig. S6: Elbow Plot, Cluster Tree, and Proportion Graph by Sex for Stromal Clusters

Suppl Table S1: a. All cluster markers. b. Cell type markers

Suppl Table S2: a. Stromal subpopulation markers. b. Marker overlap with Combes et al

Suppl Table S3: a. Downsampled cell numbers by stromal cluster by sample b. Differential gene expression output c. GO term output

Suppl Table S4. a. Monocle module genes. b. Number of cells expressing modules

## ACKNOWLEDGMENTS

This work was supported by the Northwestern University Robert H Lurie Comprehensive Cancer Center Core Facilities, including Flow Cytometry Core, Mouse Histology and Phenotyping Laboratory (MHPL) and NUSeq Core. Microscopy was provided by the Center for Advanced Microscopy of Northwetsern University and by the Microscopy and Histology Group at Stanley Manne Children’s Research Institute affiliated with Ann and Robert H. Lurie Children’s Hospital of Chicago.

We thank Dr. Matthew Schipma and Ms. Wai at NUSeq, Mr. Mehl at flow cytometry, Ms. Acar at MHPL for technical supports. We also thank Dr. Benjamin Thomson and other members of the Quaggin lab for helpful discussions.

This research was supported in part through the computational resources and staff contributions provided by the Genomics Compute Cluster which is jointly supported by the Feinberg School of Medicine, the Center for Genetic Medicine, and Feinberg’s Department of Biochemistry and Molecular Genetics, the Office of the Provost, the Office for Research, and Northwestern Information Technology. The Genomics Compute Cluster is part of Quest, Northwestern University’s high performance computing facility, with the purpose to advance research in genomics.

With gratitude to the Zell Family Foundation.

---

**Supplemental Figure S1.**
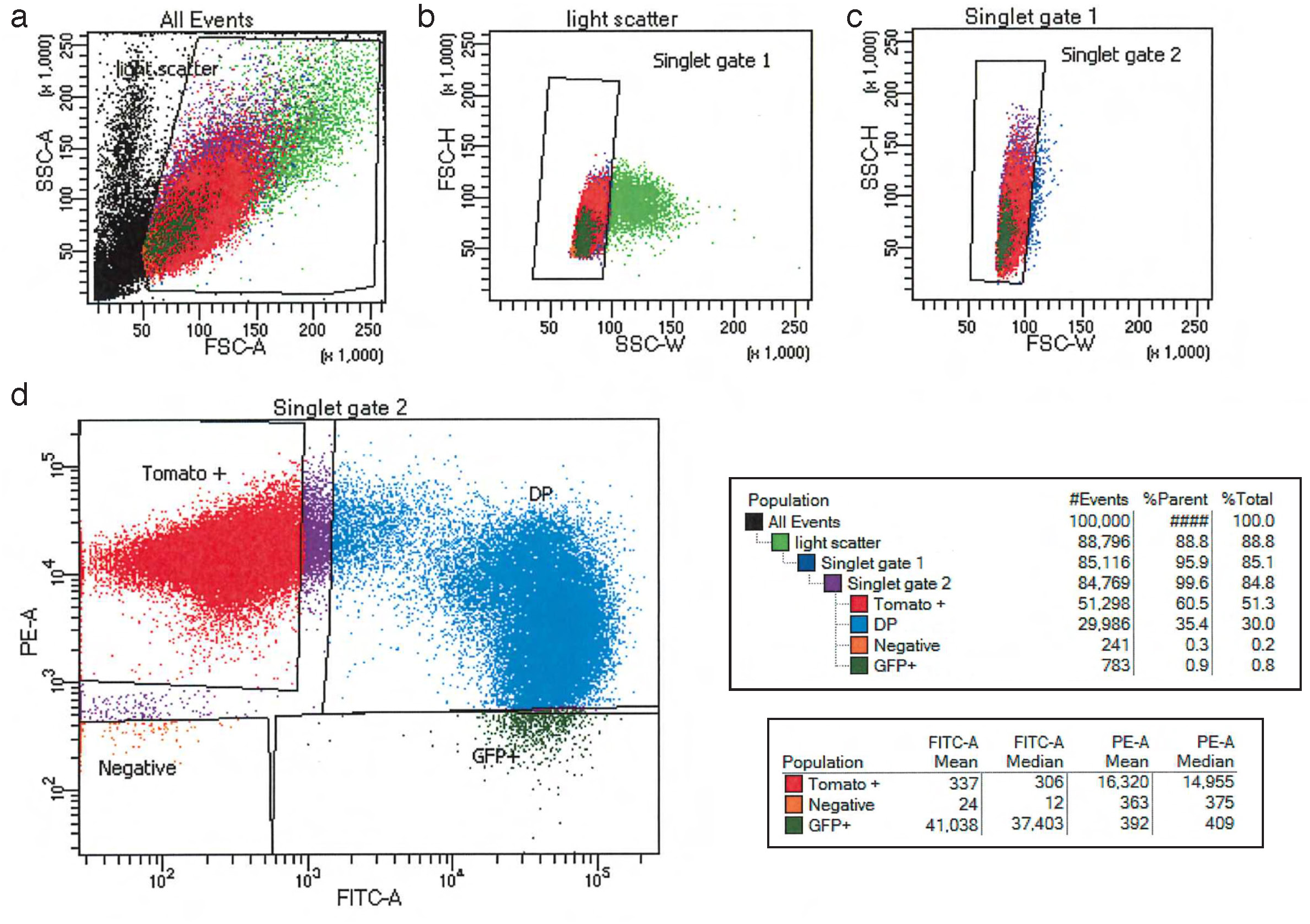
Flow cytometry analysis of single-cell suspensions from E14.5 mouse kidneys for single-cell RNA sequencing. (a) Scatter plot showing gating on cells and excluding cell debris based on forward scatter (FSC-A) and side scatter (SSC-A) properties. (b, c) Gating on single cells to exclude cell clumps. (d) Fluorescence intensity dot plot with the x-axis representing GFP fluorescence and the y-axis representing tdTomato fluorescence, illustrating gating parameters used to sort dual-positive cells from E14.5 mouse kidneys of control and *Tcf21-cKO* samples.

**Supplemental Figure S2.**
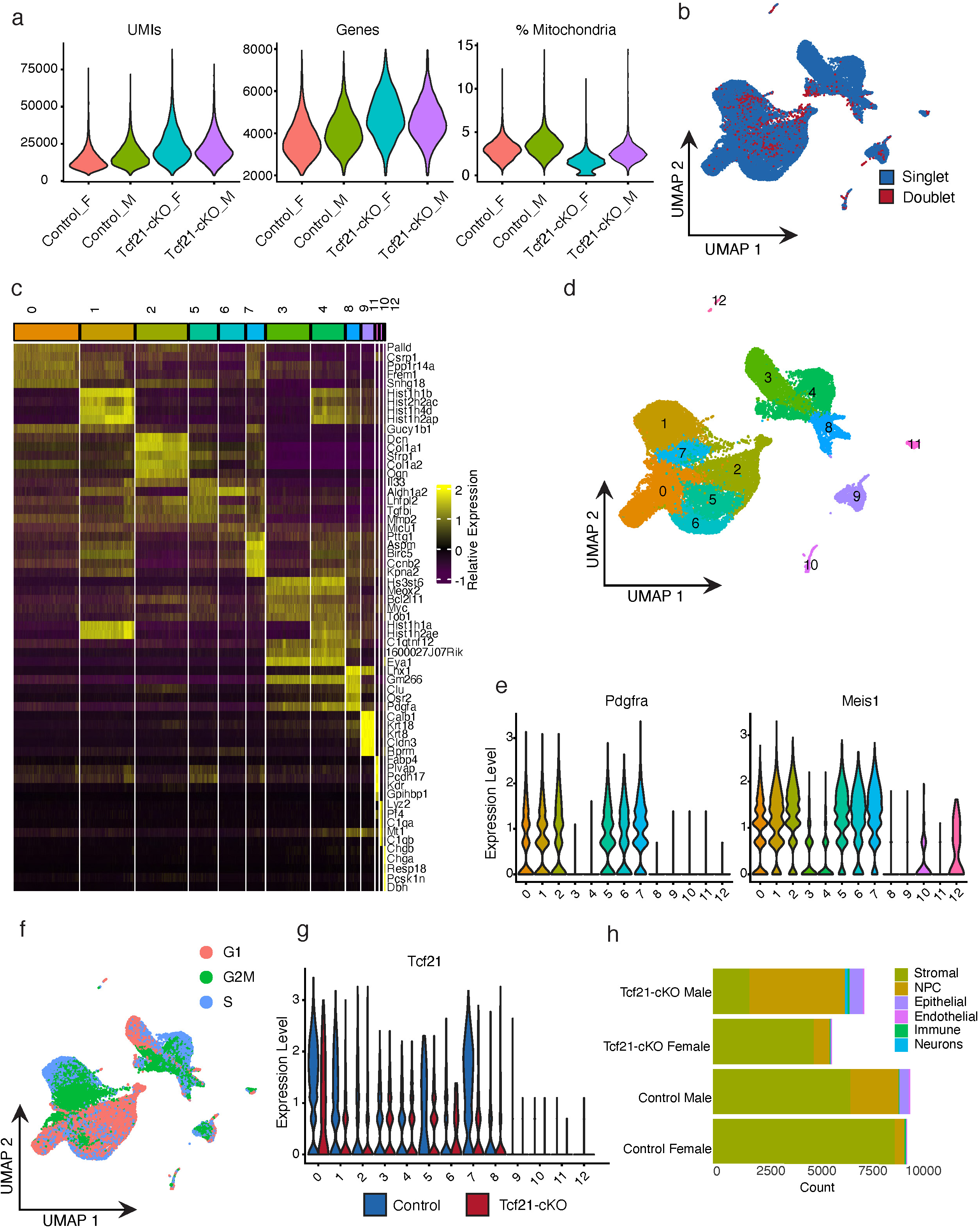
Single-cell RNA-seq analysis of Foxd1 GFP+ tdTomato+ sorted cells from control and *Tcf21-cKO* kidneys. (a) Violin plots showing number of unique molecular identifiers (UMIs), number of genes, and percentage of mitochondrial genes from control and *Tcf21-cKO* E14.5 kidney samples. (b) UMAP plot showing the singlets and doublets assignment in control and *Tcf21-cKO* mice. Doublets were removed from further analysis. (c) Heatmap depicting the relative expression of the top 5 *de novo* marker genes per cluster. Column color coding is consistent with **Suppl Fig. S2d.** (d) UMAP plot showing the unbiased clustering of cells from control and *Tcf21-cKO* mice. Thirteen clusters were defined and annotated as stromal (clusters 0,1,2,5,6,7), nephron progenitor cells (NPC) (clusters 3,4,8), epithelial (cluster 9), endothelial (cluster 11), immune (cluster 10), or neuron (cluster 12). (e) Violin plots showing log_2_-transformed gene expression of the pan-stromal markers *Pdgfra* and *Meis1* across clusters. (f) UMAP plot of control and *Tcf21-cKO* cells colored by cell-cycle stage estimated using CellCycleScoring. (g) Violin plot showing log_2_-transformed Tcf21 gene expression across clusters for control (blue) and *Tcf21-cKO* (red) cells. (h) Stacked bar plot displaying number of cells for main cell types separated by experimental group and sex.

**Supplemental Figure S3.**
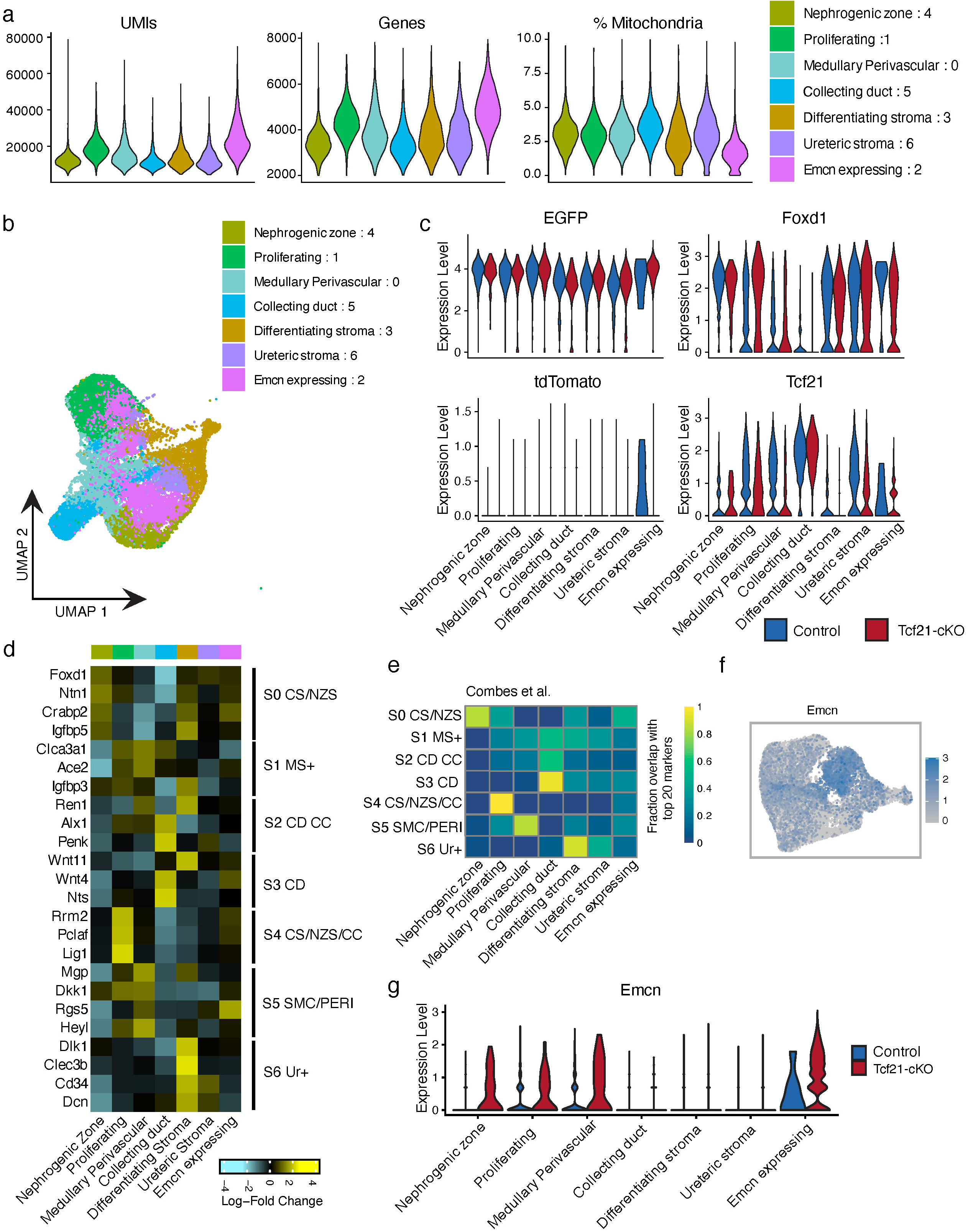
Single-cell RNA-seq analysis of stromal cells. (a) Violin plots showing number of unique molecular identifiers (UMIs), number of genes, and percentage of mitochondrial genes from control and *Tcf21-cKO* stromal subpopulations. (b) UMAP plot showing the stromal subpopulation annotations on the UMAP from **Fig. 1**. (c) Violin plot showing log_2_-transformed gene expression of EGFP, Foxd1, tdTomato, and Tcf21 across stromal subpopulations for control (blue) and *Tcf21-cKO* (red) samples. (d) Heatmap depicting the expression of stromal gene markers from Combes et al. ^26^ across our stromal subpopulations. Color scale represents log_2_ fold change in expression in the given subpopulation compared with all others. (e) Heatmap depicting the proportion of Combes et al ^26^ top 20 stromal gene markers that overlap *de novo* markers for each of our subpopulations. (f) Feature plot of log_2_-transformed gene expression of Emcn across stromal cells. (g) Violin plot showing log_2_-transformed Emcn gene expression across stromal subpopulations for control (blue) and *Tcf21-cKO* (red) cells. *(S0 CS/NZS - cortical/nephrogenic zone stroma, S1 MS+ - medullary stroma+, S2 CD CC - collecting duct-associated stroma cell cycle, S3 CD - collecting duct-associated stroma, S4 CS/NZS/CC - cortical/nephrogenic zone stroma, and cell cycle stroma, S5 SMC/PERI - smooth muscle cell/pericyte-like, S6 Ur+ - ureteric stroma.)*

**Supplemental Figure S4.**
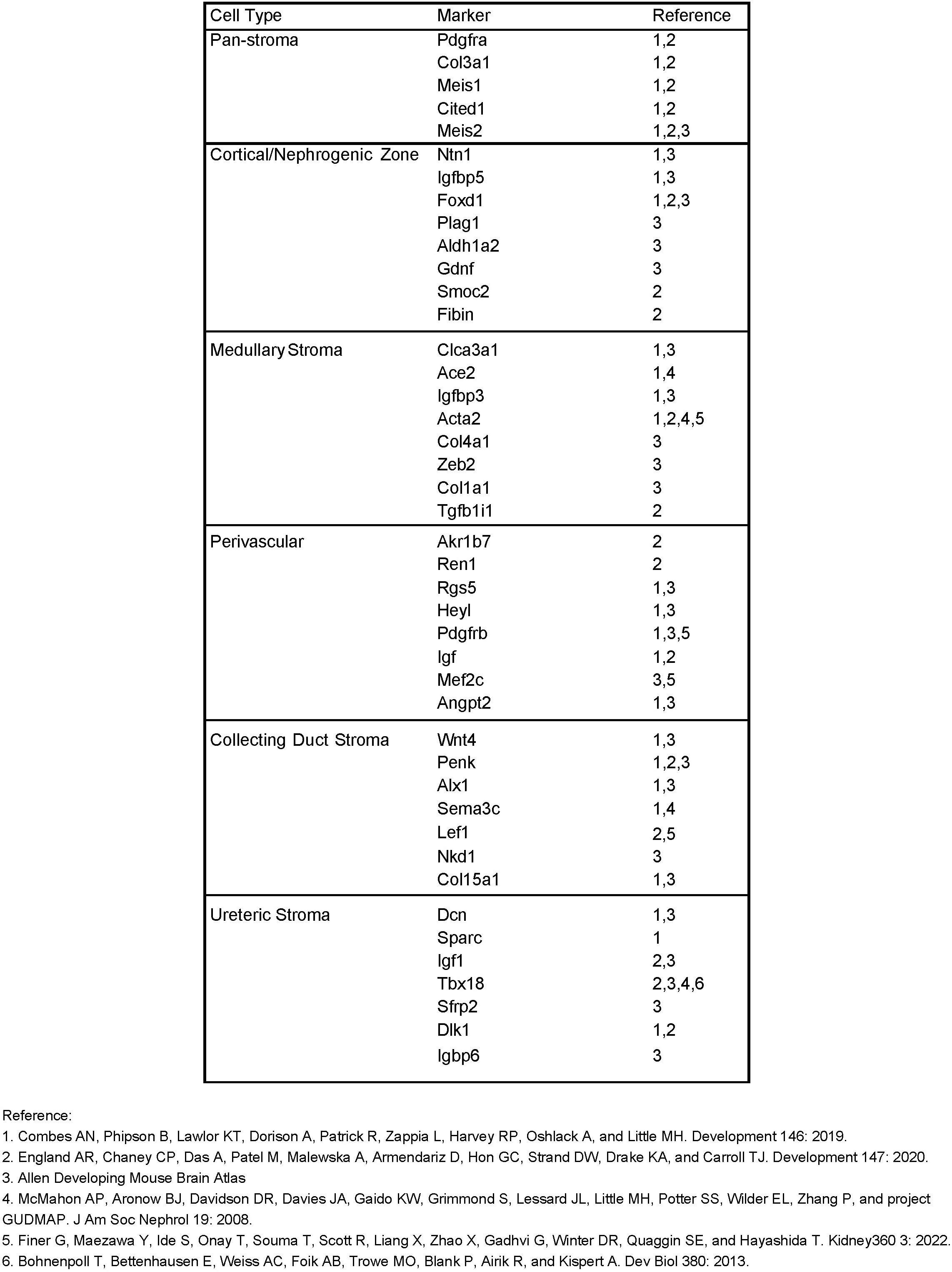
Stromal subpopulation markers for module scores. List of markers for pan-stroma and six distinct populations based on the literature references. These were used to calculate module scores for **Fig. 2c**.

**Supplemental Figure S5.**
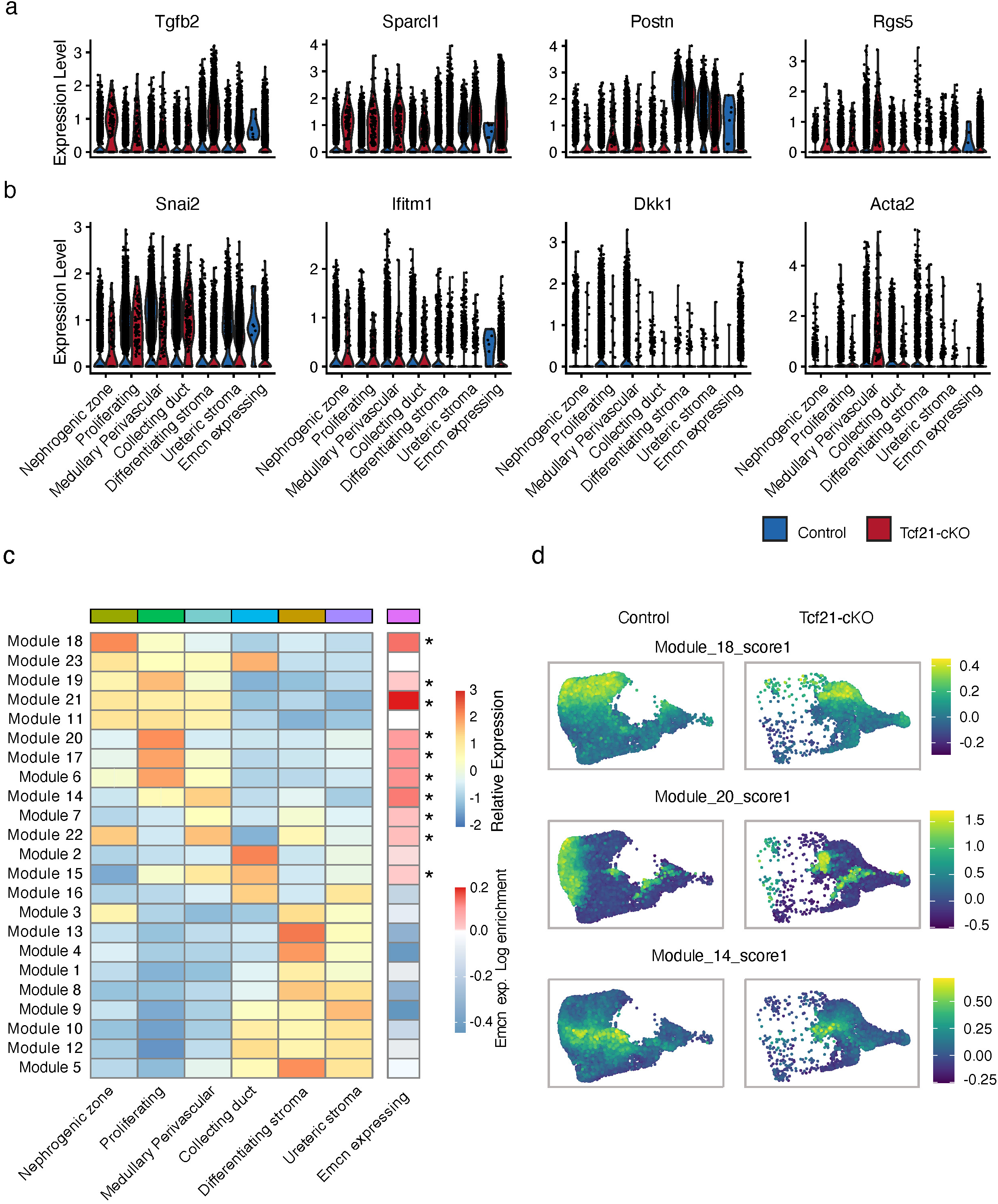
Comparison of stromal cells between *Tcf21-cKO* and controls. (a) Violin plots showing log_2_-transformed gene expression of up-regulated genes (Tgfb2, Sparcl1, Postn, Rgs5) across stromal subpopulations for control (blue) and *Tcf21-cKO* (red) cells. (b) Violin plots showing log_2_-transformed gene expression of down-regulated genes (Snai2, Ifitm1, Dkk1, Acta2) across stromal subpopulations, for control (blue) and *Tcf21-cKO* (red) samples. (c) Heatmap depicting the relative expression of 23 trajectory-based modules of co-regulated genes across control cells in each stromal subpopulation. On the right is shown the log_2_ enrichment of cells expressing these modules in the Emcn-expressing cluster compared with the rest of *Tcf21-cKO* stromal cells. * indicates p-value<0.05 by hypergeometic test. (d) Feature plot showing module scores across stromal cells split by control (left) and *Tcf21-cKO* (right) for module18 (nephrogenic zone), module20 (proliferating), and module14 (medullary/perivascular).

**Supplemental Figure S6.**
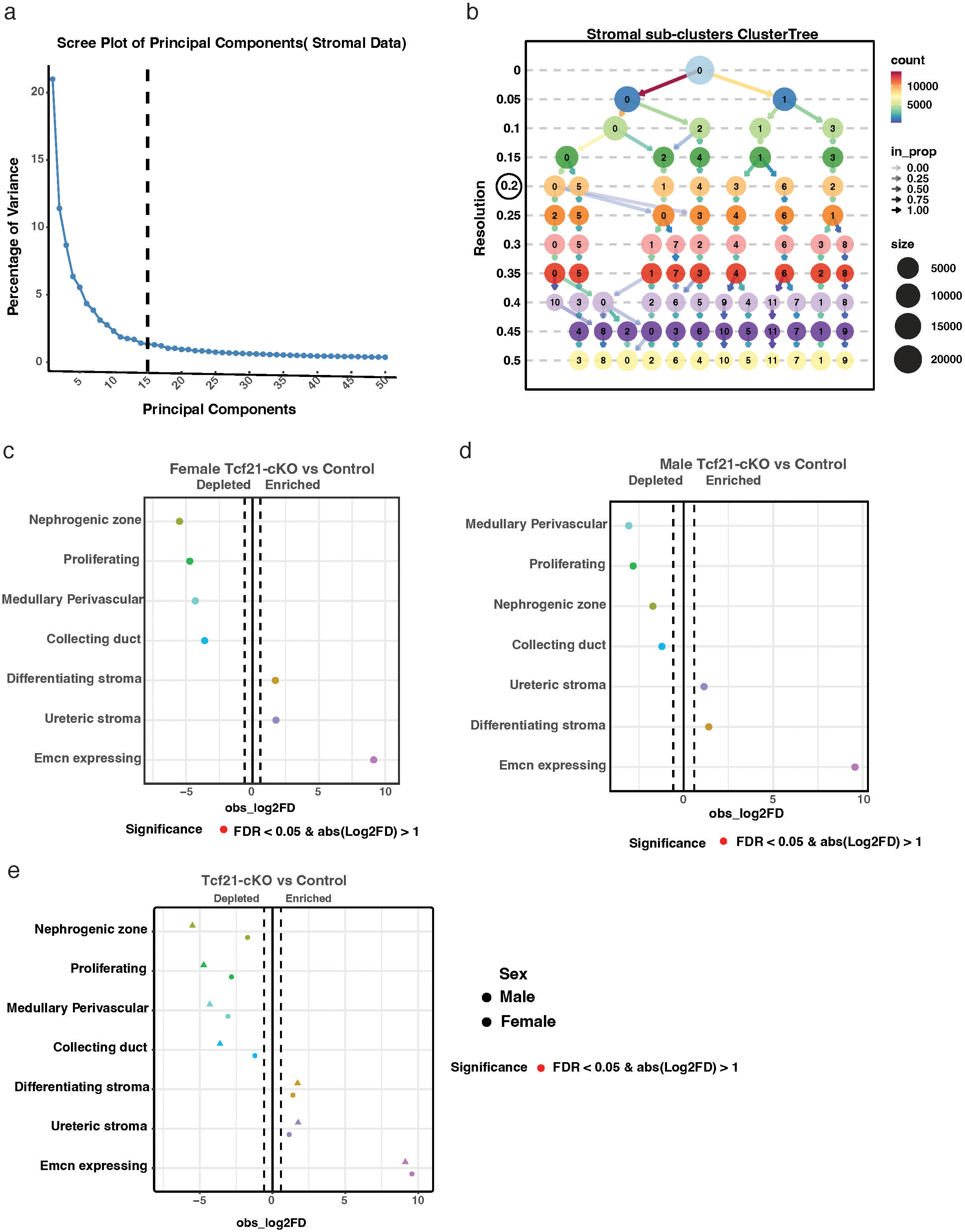
Elbow Plot, Cluster Tree, and Proportion Graph by Sex for Stromal Clusters. (a) Elbow plot depicting the optimal number of stromal sub-clusters. (b) Cluster tree illustrating robustness of stromal clusters across various resolutions. (c-e) Relative differences in cell proportions for each cluster between the *Tcf21-cKOs* versus controls in male and female samples independently. Dashes vertical lines mark absolute Log_2_ fold change (FC) > 1 compared with the control. All changes in proportion were significant after FDR was applied (scproportion permutation test; *n* = 1777 cells/group).

